# Gametophyte genome activation occurs at pollen mitosis I in maize

**DOI:** 10.1101/2021.07.26.453871

**Authors:** Brad Nelms, Virginia Walbot

**Affiliations:** Department of Biology, Stanford University, Stanford, CA 94305, USA; Department of Plant Biology, University of Georgia, Athens, GA 30602, USA

## Abstract

Flowering plants alternate between multicellular haploid (gametophyte) and diploid (sporophyte) generations. One consequence of this life cycle is that plants face substantial selection during the haploid phase^1–3^. Pollen actively transcribes its haploid genome^4^, providing phenotypic diversity even among pollen grains from a single plant. Currently, the timing that pollen precursors first establish this independence is unclear. Starting with an endowment of transcripts from the diploid parent, when do haploid cells generated by meiosis begin to express genes? Here, we follow the shift to haploid expression in maize pollen using allele-specific RNA-sequencing (RNA-Seq) of single pollen precursors. We observe widespread biallelic expression for 11 days after meiosis, indicating that transcripts synthesized by the diploid sporophyte persist long into the haploid phase. Subsequently, there was a rapid and global conversion to monoallelic expression at pollen mitosis I (PMI), driven by active new transcription from the haploid genome. Genes expressed during the haploid phase showed reduced rates of nonsynonymous relative to synonymous substitutions (d_n_/d_s_) if they were expressed after PMI, but not before, consistent with purifying selection acting on the haploid gametophyte. This work establishes the timing with which haploid selection may act in pollen and provides a detailed time-course of gene expression during pollen development.

---

Plants do not make gametes directly after meiosis, instead forming a multi-cellular haploid organism called the gametophyte. While the size of the gametophyte is reduced in flowering plants (2-3 cells for male pollen and 4-15 cells for the female embryo sac), the haploid generation retains a high degree of independence. Gametophytes actively transcribe genes, with over 60% of the genome expressed post-meiotically in pollen^4^. Many genes are required during the haploid phase, as even modest chromosome deletions are not transmitted^5, 6^. Furthermore, mutants that cannot progress through the haploid stage are routinely isolated in plant genetic screens, with hundreds of gametophytic mutants identified in *Arabidopsis* alone^7^. This widespread haploid expression exposes a large portion of the genome to natural selection in the gametophyte. Pollen, in particular, has a high capacity for selection because of the large population sizes (e.g. >10^6^ pollen grains per maize plant) and intense competition during dispersal and fertilization. It is thus not surprising that pollen selection has diverse consequences^3^: reducing inbreeding depression^8^, increasing offspring fitness^9^, and contributing to sex chromosome evolution^10^ and sex-specific differences in recombination rates^11^. Pollen selection has further been employed in breeding programs to derive cold-tolerant crop varieties^12, 13^.

The broad impacts of haploid selection in plants raises an important question: when does the haploid genome take over from its diploid parent? The haploid phase of pollen development is a complex and dynamic process^14^ that in maize lasts 20 days^15^ (Fig. 1a), roughly one third of the total time from seed to anthesis. There is no guarantee that gene products will be derived from the haploid genome immediately after meiosis. By comparison, the maternal genome controls most early events in animal (and likely plant) post-fertilization development, followed by a maternal-to-zygotic transition in which degradation of maternal products is coordinated with zygotic genome activation^16^. Does an analogous parent-to-offspring transition occur in pollen? If plants provision some portion of pollen development with diploid products, it would constrain the amount of haploid selection they experience. Here, we obtain allele-specific RNA-sequencing data from single pollen precursors across 26 days of development – from the beginning of meiosis through pollen shed. These data allow us to follow, throughout time and on a gene-by-gene basis, when expression shifts from biallelic to monoallelic during pollen development.

**Figure 1.**
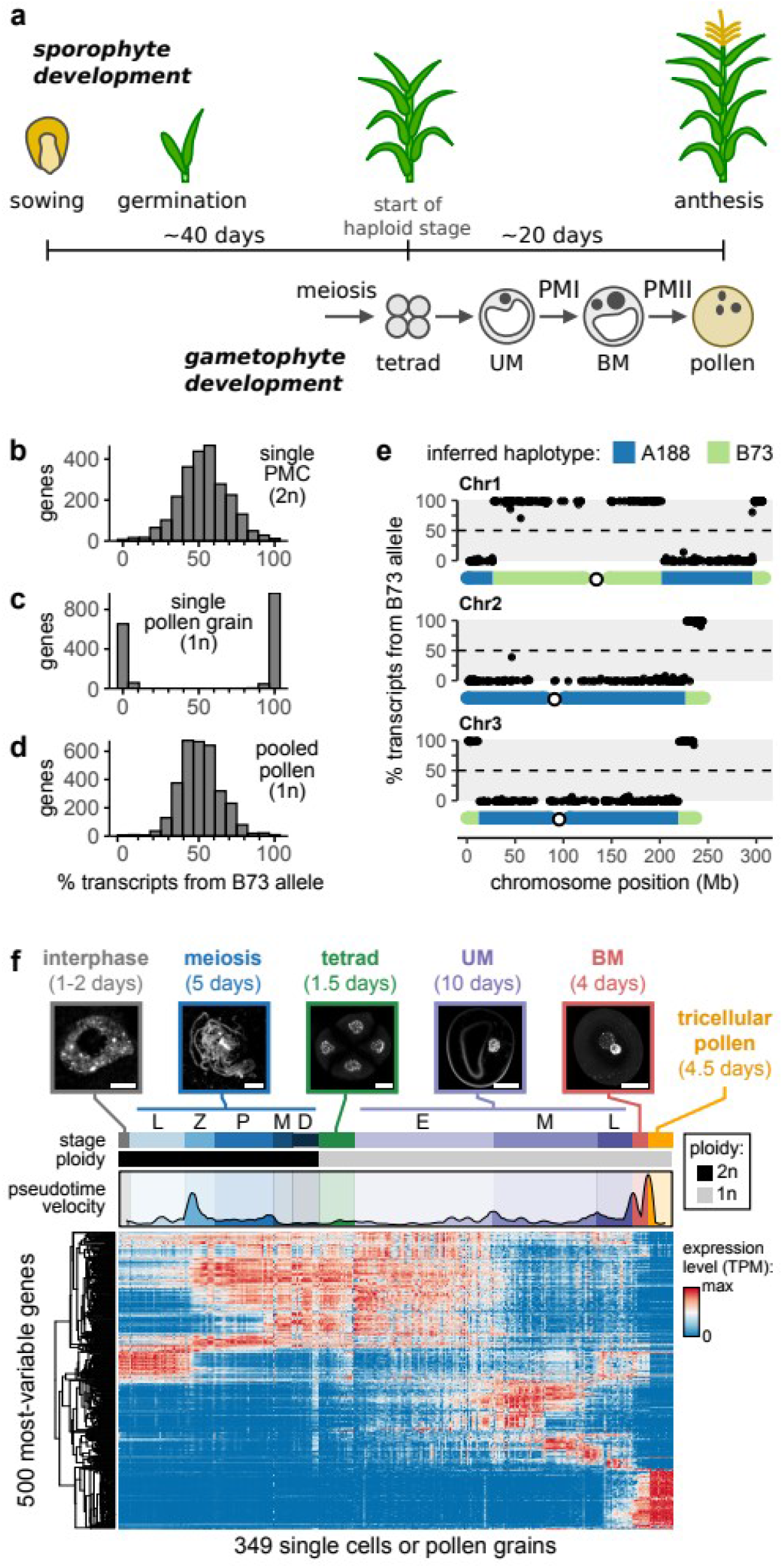
Allele-specific RNA-sequencing of single pollen precursors. (**a**) Timeline of sporophyte and (male) gametophyte development. UM, unicellular microspore; BM, bicellular microspore; PMI/PMI, pollen mitosis I and II. (**b-d**) Histogram of the fraction of transcripts matching the B73 allele for genes in (**b**) a single diploid pollen mother cell (PMC), (**c**) a single pollen grain, and (**d**) the average across pollen grains (computationally “pooled” data). (**e**) Allelic bias of genes correlates with their genomic location for the first three chromosomes of a single pollen grain. Inferred haplotype is shown below. In c-f, all genes with at least 10 genoinformative transcripts are shown. (**f**) Single-cells (UM stage and earlier) and single gametophytes (BM stage and later) were isolated from maize anthers for RNA-sequencing. Top, examplar microscopy images of paired material used for sample staging. Middle, pseudotime velocity, which quantifies the rate of expression change over time^19^; peaks in pseudotime velocity indicate periods of rapid gene expression change. Bottom, heatmap of gene expression for the top 500 most-variable genes. Scale bars are 5 μg/mL Hoechst 33342 (Sigma-Aldrich). Anthers were then mechanicallym for interphase and meiosis and 20 μg/mL Hoechst 33342 (Sigma-Aldrich). Anthers were then mechanicallym for later stages. Substages of meiosis: L, leptotene; Z, zygotene; P, pachytene; M, meiosis I division; D, dyad. UM substages: E, early; M, middle; L, late.

## Allele-specific RNA-seq of single pollen precursors

To test our ability to separate the contributions of parent (sporophyte) and offspring (gametophyte) to the transcriptome of individual pollen precursors, we first isolated single diploid pollen mother cells (PMCs; cells poised to initiate meiosis) and haploid pollen grains from an F1 hybrid between the A188 and B73 inbred lines. These two stages are separated by 26 days and represent the extremes of fully diploid expression to maximally haploid-derived expression. We detected a mean of 364,003 transcripts per sample. On average, 32.4% of transcripts could be unambiguously attributed to either the A188 or B73 alleles – hereafter referred to as genoinformative transcripts. At least one genoinformative transcript was detected for 64.3% of expressed genes.

In single PMCs, most genes were expressed from both alleles (Fig. 1b), as expected for diploid genome expression. In mature pollen grains, in contrast, genes were expressed almost exclusively from one allele (Fig. 1c). While multiple biological mechanisms can produce monoallelic expression, two pieces of evidence confirm that pollen monoallelic expression reflects expression from the haploid genome. First, there was no bias towards either the A188 or B73 alleles (Fig. 1d), as would be predicted by parental imprinting or inbred-specific effects such as presence/absence variation. Second, extensive blocks of linked genes on chromosome arms were expressed from the same parental allele, with infrequent shifts to the alternate parental allele characteristic of meiotic recombination (Fig. 1e and S1). Using the allele-specific expression data, we infer an average of 1.36 crossovers per chromosome (Fig. S2a) with more frequent crossovers towards the telomeres (Fig. S2b), in agreement with the established crossover frequency^17^ and distribution^18^ in maize. Thus, we conclude that RNA-seq of individual cells and pollen grains can distinguish expression originating from the diploid and haploid genomes.

## Gene expression during pollen development

We next profiled 349 single pollen precursors collected from 67 staged anthers, with dense sampling between pre-meiotic interphase through mature pollen (Fig. 1f and Table S1). To facilitate sample staging, precursors were collected from one anther for RNA-seq while the remaining two anthers from the same floret were fixed for microscopy. There was reproducible correspondence between gene expression and microscopic stage (Fig. 1f). As we will be comparing bi- and tri-cellular stages of pollen development with earlier unicellular stages, we collectively refer to these samples as single pollen precursors rather than single cells.

Gene expression did not change uniformly during development, but rather showed periods of rapid change interspersed with periods of relative stasis. There was a large shift in gene expression during early meiotic prophase I (Fig. 1f, arrow), consistent with an early prophase transcriptome rearrangement we described previously^19^ (Fig. S3). This was followed by several smaller waves of expression change during the rest of prophase I, a remarkably static transcriptome from metaphase I through early unicellular microspores (UMs), and another large shift in expression between UMs and bicellular microspores (BMs). We find distinct temporal expression patterns for many gene categories (Table S2-S3), including transcription factors, genes involved in meiotic recombination and synapsis (Fig. S4), and phased small RNA precursors (Fig. S5). This dataset provides an extensive time course of gene expression throughout meiosis and pollen development.

## Timing and extent of haploid expression

To follow the shift from diploid to haploid expression, we first compared the proportion of genes with biallelic and monoallelic expression in individual precursors at each stage (Fig. 2a). Genes were categorized as expressed monoallelically if they had >80% of transcripts from a single allele and as biallelically otherwise. We observed biallelic expression for the majority of genes during meiosis I (median of 83.5% biallelic genes per cell; Fig. 2a), while the cells were still diploid. Surprisingly, cells at the haploid tetrad and UM stages continued to display a similar level of biallelic expression, with a median of 82.5% biallelic-expressed genes per cell (interquartile range: 79.6% to 84.5%). Thus, pre-meiotic (biallelic) transcripts persist until the end of the UM stage, 11 days after meiosis. Subsequently, there was a rapid conversion to monoallelic expression around pollen mitosis I (PMI), with a median of 99.1% and 99.5% monoallelic-expressed genes in BMs and pollen grains, respectively. Linked genes were consistently expressed from the same allele in BMs and pollen but not earlier stages (Fig. 2a, right; Fig. S6), a characteristic sign of expression from the haploid genome. We conclude that the haploid microspore is heavily provisioned with sporophytic transcripts, followed by a sharp transition to gametophytic expression around PMI.

**Figure 2.**
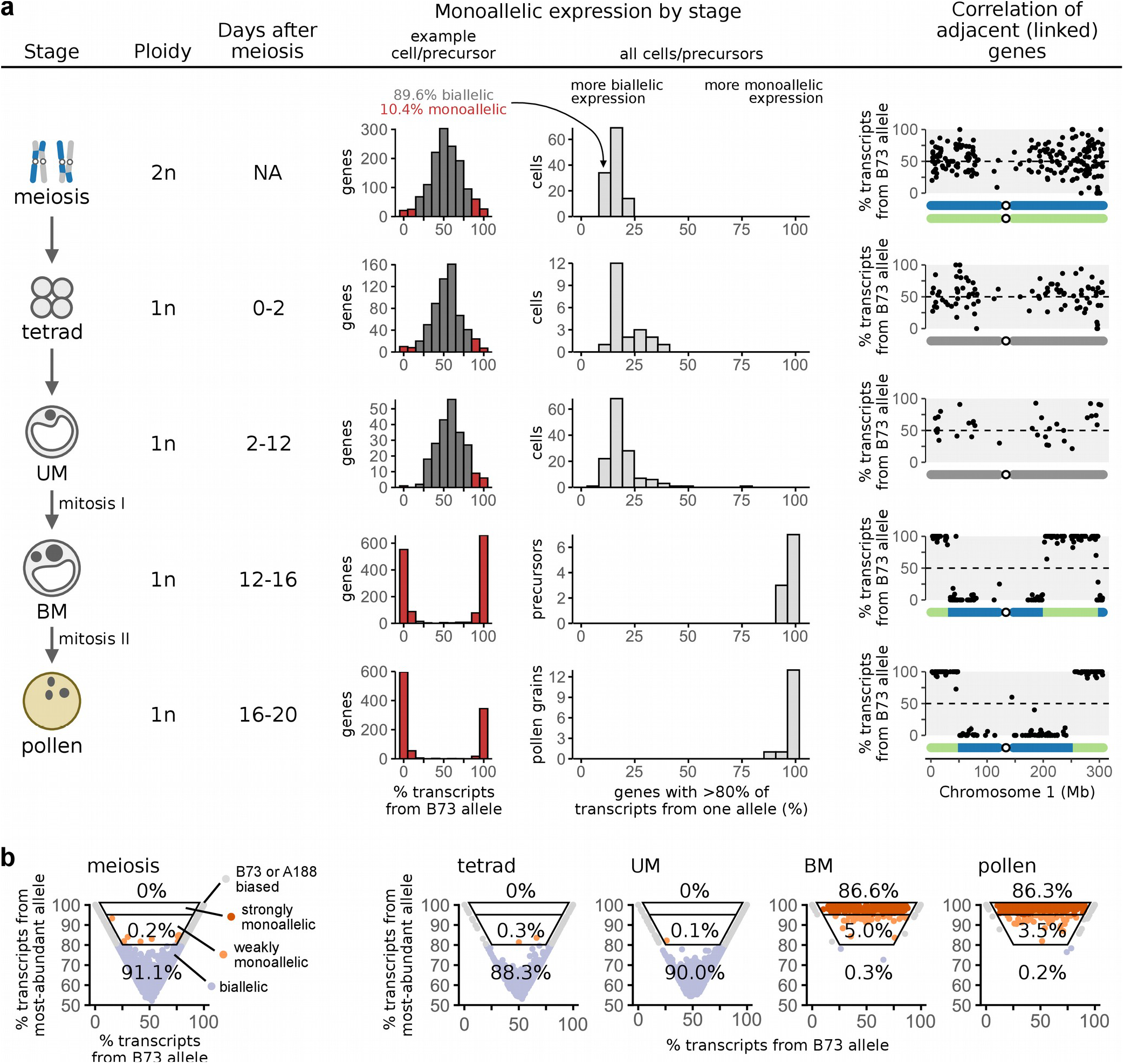
Timing of haploid expression during pollen development. (**a**) Table showing the proportion of monoallelic expression for each stage in pollen development. Column 3: mean days after meiosis when each stage begins and ends^15^. Column 4: histogram of genes, showing the fraction of transcripts matching the B73 allele in a representative precursor. Column 5: histogram of cells/precursors, showing the % of monoallelic genes in all precursors at a given stage. Column 6: % transcripts matching the B73 allele for each gene, by chromosome location. (**b**) Scatter plots showing the mean % of transcripts matching the B73 allele vs the mean % matching the most-abundant allele within a precursor for each gene, by stage. The top two boxed regions highlight genes with strong monoallelic expression (>95% from the most-abundant allele) and moderate monoallelic expression (80%-95% from the most-abundant allele), excluding genes with a consistent bias towards a specific parental allele.

While most genes had biallelic expression through PMI, does a gene cohort exist with earlier expression from the haploid genome? To answer this, we needed to distinguish haploid expression from other causes of monoallelic expression for individual genes. One unique characteristic of haploid expression is that it does not produce any bias towards a specific allele; haploid-expressed transcripts will match the A188 allele in some precursors, but the B73 allele in others, depending on the precursor haplotype. Most other causes of monoallelic expression, in contrast, result in a consistent skew towards one allele. For instance, in diploid meiotic cells 5.5% of genes were expressed monoallelically (>80% of transcripts from the most-abundant allele); however, such genes were consistently biased towards either the B73 or A188 alleles and thus can be distinguished from haploid expression (Fig. 2b). In UMs, 90.0% of genes had biallelic expression and only 0.1% had monoallelic expression (the remaining 9.9% were B73- or A188-biased). In the following stage (BMs) the reverse was true: 0.3% of genes had biallelic expression and 93.3% of genes had monoallelic expression. Thus, the shift to haploid expression is largely all-or-none: we find no evidence for genes that are expressed from the haploid genome prior to PMI or, conversely, that persist as biallelic transcripts beyond PMI. There may be early haploid expressed genes we did not sample here, as only 1068 genes had a sufficient number of genoinformative transcripts in the UM stage to make an inference about haploid expression; however, any such genes would be rare exceptions.

## Conservation of gametophyte-expressed genes

In many plant species, genes expressed in mature pollen show evidence for increased selection (both purifying and adaptive) compared to genomic background^20, 21^. One proposed explanation is that selection may be more efficient on the haploid generation^20, 21^. As our data show that the haploid genome becomes active primarily after PMI – midway through pollen development – we asked if there were differences in the average rate of nonsynonymous to synonymous substitutions (d_n_/d_s_) in genes expressed at different times in pollen development. We focused on genes with moderate transcript levels at each stage (≥100 TPM) because there was a non-monotonic relationship between expression level and d_n_/d_s_ at low levels of expression (Fig. S7), complicating the interpretation for low abundance transcripts. Genes with moderate expression after meiosis but not after PMI (i.e. genes expressed in the tetrad or UM stages but not later) showed a similar distribution of d_n_/d_s_ compared to the genomic background (Fig. 3a and S8). In contrast, genes expressed after PMI had a 30.7% lower median d_n_/d_s_, consistent with purifying selection acting in the haploid gametophyte. This stage-dependent change in dn/ds may be explained by the provisioning of haploid pollen precursors with diploid transcripts, eliminating heritable phenotypic variation until after PMI.

**Figure 3.**
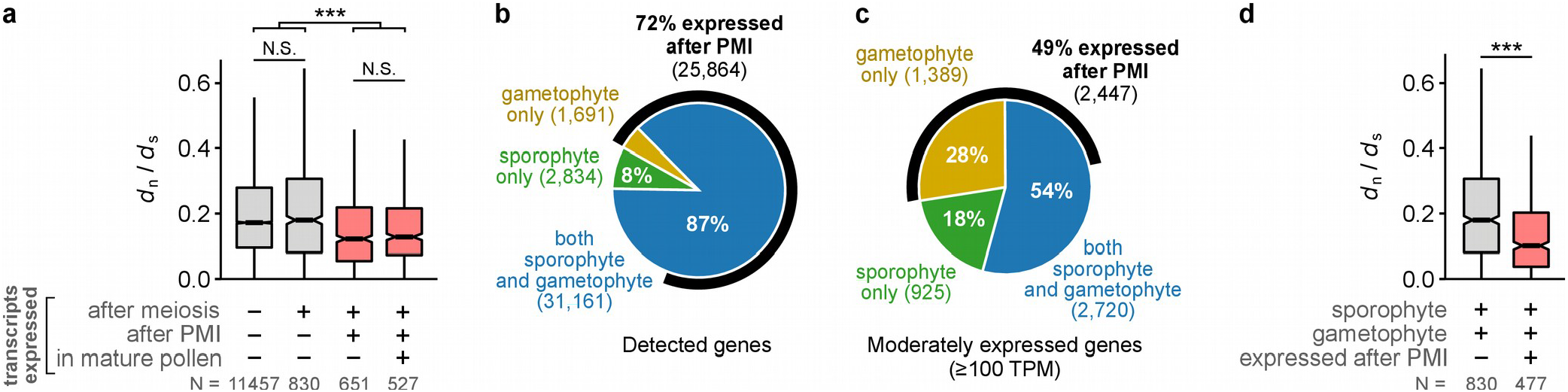
Conservation of gametophyte-expressed genes. (**a**) The ratio of the number of nonsynonymous substitutions per nonsynonymous site (*d*_n_) to the number of synonymous substitutions per synonymous site (*d*_s_) for genes expressed at different times in pollen development. Gene categories expressed after PMI are highlighted in red. (**b,c**) Proportion of genes detected (b) or moderately expressed (c) in the sporophyte, gametophyte, or both. The number of genes expressed after PMI are also indicated. (**d**) *d_n_/d_s_* for genes expressed in both the gametophyte and sporophyte stages, separated based on whether they were expressed after PMI. For (a) and (d), only moderately expressed genes (≥100 TPM) were considered. Boxplots show the median (horizontal line), interquartile range (shaded area; IQR), and whiskers extending up to 1.5 x IQR. Gene categories expressed after PMI are shaded red. ***, p < 0.001, Wilcoxon test adjusted for multiple hypothesis testing with Holm’s method.

We next estimated the fraction of genes expressed in the diploid sporophyte that might be subject to haploid selection in pollen. To identify sporophyte-expressed genes, we obtained expression data from whole seedlings (roots and shoots), defining sporophytic genes as those expressed in either seedlings or diploid pollen precursors. Consistent with prior results^4, 22, 23^, we found that a large fraction of the genome is expressed during both diploid and haploid stages: 87.3% of genes had detectable transcripts in both the sporophyte and gametophyte (Fig. 3b) and 54.0% were moderately expressed in both (≥100 TPM; Fig. 3c). Of these, a substantial portion were expressed after PMI and thus potentially subject to haploid selection (Fig. 3b,c); this subset had a significantly lower median d_n_/d_s_ (Fig. 3d). In total, 25,864 genes were detected and 2,447 were moderately expressed after PMI.

## Widespread gametophyte genome activation at PMI

What is the contribution of new transcription vs transcript turnover to the shift to haploid expression? RNA dynamics usually cannot be inferred from steady-state transcript levels alone, because opposing changes in the rate of RNA synthesis and degradation can produce similar effects on transcript abundance^24^. Our data provide a way to separate synthesis from degradation, however, because during the haploid phase any new transcription can only come from one allele. We find that the mean number of transcripts per precursor changed substantially during pollen development (Fig. 4a), suggesting large differences in the relative rate of new synthesis vs degradation between stages. There was a steady decrease in transcripts per cell from the peak during early meiosis to the minimum at the UM stage. This was followed by a sharp, 7.5- fold increase in the total number of transcripts per precursor between late UMs and BMs (95% confidence interval (CI) = 3.0 to 14.2-fold; bootstrap test), indicating that substantial new transcription may drive the shift to monoallelic expression during this period. Indeed, 7,361 genes had a ≥2-fold increase in absolute transcript abundance between late UMs and BMs (Fig. 4b), with this increase attributable to the more-abundant (haploid) allele (Fig. 4c). In contrast, the less-abundant allele remained relatively constant between UMs and BMs (median fold change of 0.02; Fig. 4d). This suggests that premeiotic (biallelic) transcripts continue to persist into the BM stage for many genes, but that a large increase in new transcription overtakes pre-existing transcript levels to produce a net shift towards monoallelic expression. We conclude that the transition to haploid expression is driven by new transcription and gametophyte genome activation, with degradation of sporophytic transcripts playing a relatively minor role at the transition.

**Figure 4.**
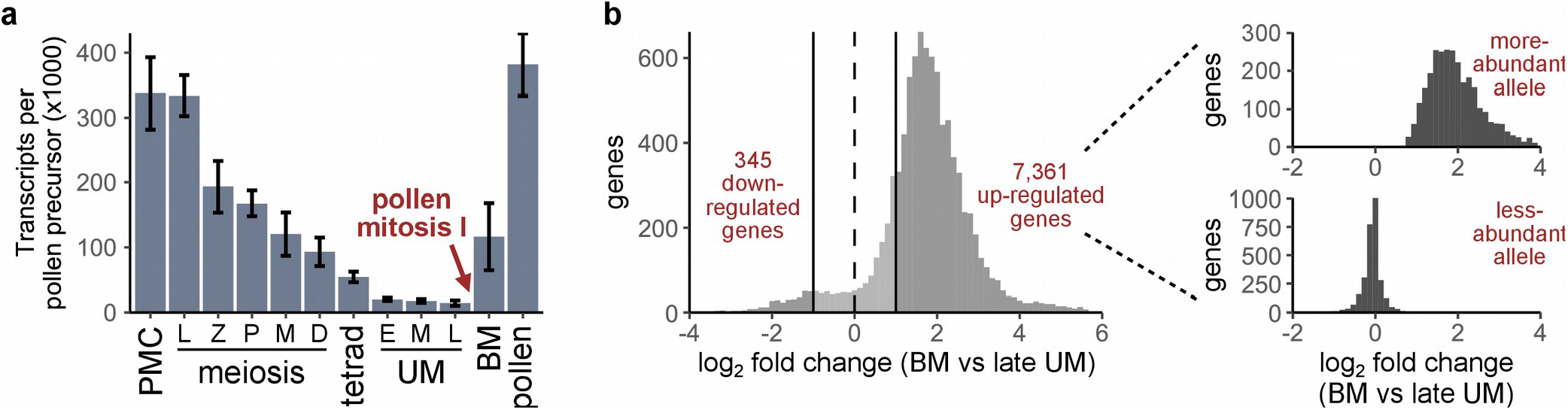
Widespread gametophyte genome activation at pollen mitosis I. (**a**) Total transcripts detected per pollen precursor, by stage. Shown are trimmed means (trim = 0.2) ± standard errors, estimated by bootstrapping. (**b**) Left, log_2_ fold change in absolute transcript abundance between the late UM and BM stages for genes with a mean expression level ≥1 transcript per precursor. Right, log_2_ fold change in transcript abundance for transcripts mapping to the more- and less-abundant allele (top and bottom, respectively), showing only genes with a 2-fold or greater increase in overall transcript abundance. Up-regulated genes show an increase in transcript levels for the more-abundant (haploid) allele only.

*De novo* motif analysis identified the RY repeat (CATGCA[TG]) as significantly enriched in the promoters of the top 200 most upregulated genes, with 35/200 promoters (17.5%) having a perfect match to the full RY repeat (6.1-fold enrichment; p = 7.1x10^-15^, Fisher’s exact test) and 72 (36%) containing the minimal ‘CATGCA’ motif (2.4-fold enrichment; p = 6.2x10^-9^, Fisher’s exact test). The RY repeat is the binding site for three paralogous transcription factors that regulate embryogenesis in *Arabidopsis*^25^ *(ABI3, FUS3,* and *LEC2).* Although the RY repeat has no known function in pollen development, conserved RY repeats have been noted in the pollen-specific β-expansin genes^26^. This sequence may serve as the binding site for a transcription factor that contributes to gametophyte genome activation. We note that *ABI3* and *ABI19*, two of 4 maize orthologs of *FUS3/LEC2*, are specifically expressed in the embryo and late-stage anthers^27^.

## Gene regulation before PMI

Prior to PMI, there appeared to be very little new transcription from the haploid genome, as evident in the continued biallelic status of most transcripts (Fig. 2). However, there were still clear changes in relative transcript abundance entering the mid- and late-UM stages (Fig. 1f, Table S2-S3). To understand how the transcriptome might change in the absence of new transcription, we examined the absolute transcript abundance attributable to each allele for UM-expressed genes. Most genes showed biallelic transcript loss in UMs, ranging from rapid loss (Fig. S9a) to slower degradation over time (Fig. S9b). Thus, differences in mRNA half-life explain some expression changes during the UM stage. Surprisingly, there were also many genes with a sharp, biallelic increase in transcripts within UMs (Fig. S9c,d). What could cause a biallelic transcript increase in a haploid cell? One possibility is that these transcripts were synthesized pre-meiotically but then stored and only processed later. Our sequencing libraries enrich for polyA+ RNA and so would not detect stored RNAs with a short or missing polyA tail. The storage of un-processed RNAs has been described in other pathways such as seed development^28^, and would provide a mechanism to regulate gene expression during the UM stage without new transcription from the haploid genome. Altogether, our data show that the UM transcriptome is not static despite the lack of new transcription.

## Discussion

Here we show that diploid-derived transcripts persist long into haploid phase of maize pollen development, followed by a rapid transition to monoallelic expression around PMI. We propose to call this the sporophyte-to-gametophyte transition (SGT), in analogy to the maternal-to-zygote transition (MZT), as both represent a shift from parent-to-offspring expression between generations. The widespread provisioning of the UM with sporophytic transcripts indicates a substantial parental investment in the developing gametophyte and implies that most cellular processes are under sporophytic control for the first half of pollen development.

Why might the SGT be delayed until pollen mitosis I? One explanation is that PMI sets up the gametophyte germline (generative cell) and soma (vegetative cell). Active transcription is associated with an increased mutation rate^29^, and so limiting transcription during the UM stage might reduce transcription-coupled DNA damage and accessibility of the genome to transposons. After PMI, the somatic vegetative cell is far more transcriptionally active than the generative cell^30^ and could accommodate transcription without an associated risk to the germline. It will be important to establish whether SGT timing varies between species and between male and female gametophytes. Is PMI a conserved moment of gametophyte genome activation? Or, does the SGT occur at different times in distinct plant lineages?

The substantial increase in new transcripts around PMI suggests that the SGT is driven by gametophyte genome activation resulting in new transcription, although the mechanisms of this activation are unknown. It is unlikely that the mitotic division itself is required to activate transcription, as vegetative cell-like development continues even when PMI is blocked^31–33^, and several gametophytic mutants have been isolated that disrupt PMI^7, 23^. Our working hypothesis is that the SGT begins immediately before PMI rather than during it. Many substantial changes have been observed around PMI, including broad shifts in protein and RNA composition^34^, transposon activity (in Arabidopsis^35^), and histone modifications^36^. There is much to learn about how these pathways are coordinated to establish the independence of the gametophyte generation.

In predominantly diploid organisms, the scope of haploid selection has long been debated^1^. Plants are generally accepted to experience greater haploid selection than animals, in part because they require many genes to complete the haploid phase^5–7^. In contrast, fully enucleate animal sperm are viable and can fertilize an egg^1^. This distinction between kingdoms may be more nuanced than previously thought. A recent study found that many genes have haploid-biased expression in mammalian sperm^37^, and consequently animal sperm may have a greater amount of heritable phenotypic variation than often assumed. Here we demonstrate an absence of haploid transcript accumulation for more than half the haploid phase in maize pollen, limiting the time period that haploid selection may act in the male plant gametophyte. The ability to measure allele-specific expression directly in haploid gametes and gametophytes will provide needed clarity on this small but important stage of the life cycle.

## Methods

### Isolating single pollen precursors for RNA-sequencing

Plants were grown in a greenhouse at Stanford, CA with a 14-h 31 °C day at 50% summar solar fluence / 10-h 21 °C night. All samples were collected mid-day from an F1 hybrid between maize (*Zea mays*) inbred lines A188 and B73, with A188 as the female parent. Plants were harvested immediately before use, and anthers were dissected from the central tassel spike. Anther length was used as a preliminary staging guide, later refined by microscopy (see below).

For anthers at the tetrad stage and earlier, we found that cells could be mechanically released from fixed anthers without requiring enzymatic digestion. Fixation with 3:1 ethanol:acetic acid preserved RNA quality and was compatible with single-cell library preparation. All three anthers from a floret were fixed in 3:1 ethanol:acetic acid on ice for 2 h. One anther was withdrawn, rinsed twice in cold phosphate-buffered saline (PBS; Sigma-Aldrich P4417), cut transversely with a #11 scapel, and gently pressed to release developing meiotic cells. Single meiotic cells were identified by their large size and distinctive morphology and manually retrieved with a 33 gauge syringe needle. The cells were then washed in PBS and placed individually on the cap of an 8-tube PCR strip (Axygen Low Profile 8-Strip PCR Tubes; Fisher Scientific 14-223-505). The presence of a single meiotic cell without attached debris was confirmed microscopically during both pick-up and release (10X magnification, Nikon Diaphot). Cells in the dyad or tetrad stage often remained attached after release from the anther; they were aspirated up and down with the syringe needle to liberate individual cells prior to isolation. After isolating a set of 8 cells, the PCR caps were attached to PCR tubes, flash frozen in liquid nitrogen, and stored at -80 °C.

For anthers at the unicellular microspore stage and later, the developing pollen precursors were loosely associated with the anther wall and could be released mechanically without enzymes or fixation. Two of three anthers from a floret were fixed in 3:1 ethanol:acetic acid and stored for microscopy. The remaining (unfixed) anther was cut transversely with a #11 scapel, then gently pressed in a drop of PBS to release pollen precursors. Single precursors were aspirated with a blunted 29 gauge needle (for unicellular or bicellular microspores) or 26 gauge needle (for mature pollen), placed on the cap of an 8-tube PCR strip, and flash frozen as described above. Anther lengths were measured with a stage micrometer (Fisher Scientific) as listed in Table S1.

### Cytological staging and image acquisition

Fixed anthers were washed in ice cold PBS and placed on a glass slide in a drop of PBS with 10 μg/mL Hoechst 33342 (Sigma-Aldrich). Anthers were then mechanicallyg/mL Hoechst 33342 (Sigma-Aldrich). Anthers were then mechanically macerated with a scapel to release pollen precursors, and a #1.5 coverslip (Zeiss) was placed on top. Images were taken with a Leica SP8 confocal laser-scanning microscope using a 40X 1.2 n.a. glycerin immersion objective and a 405 nm excitation laser. Anther stage was scored based on the morphology of the Hoechst-stained chromosomes using previously defined criteria^19, 38^. All cytological staging was performed blind to sample identity (i.e. without knowledge of anther length, gene expression, or other sample information).

### Illumina library preparation

Sequencing libraries were prepared by CEL-seq2^39^ as described previously^19^ with one addition: previously, we noticed that the read distribution was uneven within these libraries, with reads often mapping to one or a few specific positions on a transcript. We suspected this occurred during reverse transcription of the amplified RNA library, a step that uses a random hexamer primer with a long overhanging sequence at the 5’ end (5’-<u>GCCTTGGCACCCGAGAATTCCA</u>NNNNNN, the overhang is underlined). This overhanging sequence may hybridize to specific areas of the transcript, leading to preferential binding at those sequences. To avoid this problem, we modified the reverse transcription primer to the following: 5’-gaautcucgggu<u>GCCTTGGCACCCGAGAATTCCA</u>NNNNNN (original overhang is underlined; newly added sequence is in lower case) The new nucleotides on the 5’ end form a hairpin with the overhang sequence, blocking this overhang from contributing to priming during reverse transcription. After the reverse transcription step, the hairpin would inhibit downstream PCR reactions and must be removed. To remove the hairpin, the library was treated with a 1:20 dilution of USER enzyme (New England Biolabs) at 37 °C for 30 min; USER cuts DNA at uracil bases, thus degrading the hairpin portion of the primer. In test libraries, this modification produced a more even size distribution of amplified products. After the modified second reverse transcription step, library preparation was continued as described^19^. Libraries were sequenced on an Illumina NovaSeq or HiSeq 4000 instrument by Novogene Corporation (Sacramento, CA) using paired-end 150 bp reads.

### Allele-specific transcript quantification

The first read of each read-pair contained a cell-specific barcode and a 10 bp unique molecular identifier^40^ (UMI), while the second read contained transcript sequence. Paired-end reads were demultiplexed based on the cell-specific barcodes, requiring a perfect match to one of the expected barcode sequences (Table S4). Then the UMIs from read 1 were extracted and appended to the read 2 sequence identifiers. The remainder of processing was performed using the demultiplexed second reads. Prior to alignment, reads were trimmed and filtered using Fastp v0.20.0^41^ with parameters -y -x -3 -f 6. This removed Illumina adapter sequences and low-quality reads (Phred <= 15), removed the first 6 bases from the 5’ end, and trimmed poly-X sequences and low quality bases from the 3’ end.

Genome sequences were obtained for the B73 reference genome^42^ (AGPv4) and A188 genome^43^. To avoid bias in aligning B73 vs A188 alleles to the genome, the B73 genome was masked by replacing single-nucleotide polymorphisms (SNPs) that differ between the A188 and B73 genomes with Ns. Reads were then aligned to the masked B73 reference genome using Hisat2 v2.1.0^44^. Mapped reads were assigned to a gene if they overlapped the annotated gene locus. After alignment, SNPsplit v0.3.2^45^ was used to analyze the previously masked SNP positions and split the reads into four categories: those matching the A188 genome, those matching the B73 genome, those lacking SNPs that could separate the two alleles (“unassigned” reads), and those with conflicting SNPs (“conflicting” reads).

For transcript counting, reads mapping to the same gene with a similar UMI were collapsed using the UMI-tools v1.0.0^46^ dedup function; this algorithm uses graph-based clustering to group reads with similar UMIs while accounting for potential sequencing errors. In cases in which multiple reads were associated with a single transcript, the allele calls for these reads were pooled as follows: if all reads were “unassigned”, then the transcript was labeled as “unassigned”; if all reads were assigned to only one of the two alleles (i.e. only A188 or only B73) or were unassigned, then the transcript was assigned to that allele (e.g. if there were three reads associated with a transcript, including two unassigned reads and one assigned to A188, then the transcript was assigned to A188); if any read was “conflicting” or if there were both A188 and B73-assigned reads, then the transcript was labeled as “conflicting”. In total, 66.0% of transcripts were unassigned, 33.4% were assigned to the A188 or B73 allele, and 0.58% were conflicting.

### Quality control

Three lanes of sequencing were obtained for this study. For all lanes, samples with under 2,000 UMIs were discarded. There were also additional quality control criteria applied to lanes 1 and 3: For lane 1, there was evidence of an Illumina index-hopping artifact where a subset of reads were misannotated to the incorrect sample during sequencing (see below and Supplementary Note 1); additional samples were excluded from this lane to reduce the impact of index hoping (Supplementary Note 1). For lane 3, there appeared to be a library synthesis problem in many of the libraries, as 68 samples (72%) were below the 2000 UMI threshold and many other samples were marginally above it. From this lane, we kept only 7 samples that had good sequence diversity (Table S1); these samples contained 87% of all reads from the lane and had a median of 155,521 transcripts per sample, compared to a median of 498 for the excluded samples. Lane 2 was obtained under conditions where the index-hopping artifact was reduced and contained good sequence diversity; no additional quality control criteria were applied to this lane.

### Illumina index-hopping

Since the introduction of patterned flow cells, Illumina instruments have suffered from an index-hopping artifact in which the multiplexed sample barcodes can be exchanged during sequencing, leading to a fraction of reads being assigned to the wrong sample^47, 48^. There was evidence of this artifact in our data, including larger than expected correlations between samples with certain barcode combinations (Supplementary Note 1) and a small but significant number of reads mapping to impossible barcode combinations (combinations of CEL-seq and Illumina barcodes that were not used in the experiment). Several steps were taken to mitigate the impact of index-hopping and ensure that our conclusions were robust to this artifact; these include correcting the data using methods to discard reads that likely arise from index-hopping^48^, excluding samples that had a higher likelihood of index-hopping artifacts, and, importantly, confirming the findings with independent RNA-sequencing data obtained under conditions where index-hopping was minimized. A further discussion of these issues is provided in Supplementary Note 1.

### Refining developmental stages using gene expression

Pollen precursors were first assigned a developmental stage by microscopy, and then ordered within a stage using ‘pseudotime’ – a dimensionality reduction technique to infer the temporal order of samples based on gene expression^49^. To calculate pseudotime, expression data were first normalized to transcripts per million (TPM). Then the data were log-transformed after adding a pseudocount of 148.4; this pseudocount corresponds to 1 transcript per cell in a cell at the 10^th^ percentile of total detected transcripts. Genes were filtered to remove any that did not have at least 10 UMIs in at least 10 pollen precursors. Next, the 2000 most variable genes were selected (as measured using the Fano factor), and the data were transformed into principal components (PCs). Finally, a principal curve was fit to the top 10 PCs using the R package princurve^50^. Pseudotime was calculated separately for cells in meiosis and for haploid pollen precursors (tetrad stage and beyond).

Periods of more rapid gene expression change were then identified using ‘pseudotime velocity’^19^. Pseudotime velocity calculates the rate of change in pseudotime of adjacent ordered cells, reaching a larger value when gene expression changes more abruptly. Peaks in pseudotime velocity were selected as stage boundaries to divide unicellular microspores into early, middle, and late substages (Fig. 1f). The heatmap of expression along with pseudotime velocity (Fig. 1f) was plotted using the R package ‘ComplexHeatmap’^51^.

### Differential gene expression

Bootstrapping was used to estimate the mean log-transformed change in expression level between consecutive developmental stages. The following six stage comparisons were made: (i) early leptotene vs pachytene, (ii) pachytene vs M1, (iii) early UM vs mid UM, (iv) mid UM vs late UM, (v) late UM vs BM, and (vi) BM vs pollen. Bootstrapping *p*-values were adjusted for multiple hypothesis testing using the Benjamini-Hochberg procedure. Genes with an average expression level of less than 1 transcript per single pollen precursor were excluded from analysis. Differential expression was calculated using both the absolute numbers of transcripts per precursor (Table S2) and TPM-normalized data (Table S3).

### Quantifying the fraction of biallelic-expressed genes

Biallelic-expressed genes were defined as those genes with under 80% of transcripts from the most-common allele. The fraction of biallelic-expressed genes in a pollen precursor was calculated using genes with at least 10 genoinformative transcripts in that precursor. Genes were excluded from these calculations if >90% of all transcripts in the entire dataset came from the same parental allele (either A188 or B73).

### Seedling RNA-seq data

To determine seedling-expressed genes, 10-day-old whole seedlings (roots and shoots) were frozen in liquid nitrogen and ground to a fine powder with a mortar and pestle. RNA was extracted using a Qiagen RNeasy Plant Mini Kit according to manufacturer instructions and sequencing libraries were prepared and analyzed as described for pollen precursors.

### Conservation of gametophyte-expressed genes

The ratio of nonsynonymous substitution per nonsynonoymous site (*dn*) to synonymous substitution per synonymous site (*ds*) was calculated for all orthologous gene pairs between maize and other grasses with the CoGe SynMap2 tool^52^ (genomevolution.org), using the masked *Zea mays* v4 genome (id52733), *Sorghum bicolor* v3 genome (id31607), *Oryza sativa* v7 genome (id16890), and *Brachypodium distachyon* v3 genome (id39836). To avoid analyzing substitutions between non-orthologous gene pairs, putative orthologs were only considered if they were reciprocal best hits and shared at least 80% nucleotide identity.

Gametophyte-expressed genes were defined as those expressed in the haploid stages of pollen development (tetrad, UM, BM, and Pollen) at an expression threshold of 0 TPM (for Fig. 3b) or 100 TPM (for Fig. 3c-d). Sporophyte genes were defined as those genes expressed in diploid pollen precursors (meiosis or pre-meiotic interphase) or in seedlings using the same expression cutoffs. At either expression cutoff, there was a subset of genes expressed exclusively in the tetrad and/or UM stages. It was ambiguous how to classify these genes, as they were found only in the haploid gametophyte stage but were most likely expressed from the diploid sporophyte genome (see Results). All genes in this category with ≥10 genoinformative transcripts had transcripts matching both alleles. In Fig. 3b-d, we categorized this group as expressed in both the sporophyte and gametophyte (but before PMI) as we suspect they were synthesized before the end of meiosis. The conclusions do not change if a different categorization is chosen.

### *De novo* motif analysis

MEME 5.3.3^53^ was used for *de novo* motif discovery in the promoters of the top 200 genes upregulated during PMI using parameters -dna -mod anr -nmotifs 10 -minw 6 - maxw 15 -objfun classic -revcomp -markov_order 0. The top 200 genes were defined as those with the greatest increase in absolute transcript abundance between late UMs and BMs with a false discovery rate cutoff of 0.01 (Table S2). Promoters were defined as the first 500 bp upstream of the transcription start site for each gene. The top three motifs were far more significant than all the others: ‘AWAAAAAAAWATAAA’ (E-value = 7.6x10^-56^), ‘CCBSCBCCBCCKCSC’ (E-value = 2.4x10^-42^), and ‘CATGCATGCA’ (E-value = 3.8x10^31^). As the first two were long polynucleotide stretches (polyA and polyC), only the RY repeat (‘CATGCATGCA’) was considered further. Separate from *de novo* motif discovery, we estimated the enrichment of the RY repeat (using the smaller established motif sequences: CATGCATG and CATGCA) in the promoters of upregulated genes compared to a background gene set of 5099 genes. The background genes were selected based on their low expression in the BM stage (< 10 TPM) and an expression level between 4 and 900 TPM in seedlings and/or pollen precursors. These expression cutoffs match the expression level to the upregulated geneset, as the upregulated genes had a mean expression between 4 and 900 TPM across pollen precursors.

### Statistical methods

Significance in the difference in *dn/ds* between stages (Fig. 3 and S8) was calculated with a two-sided Wilcoxon signed-rank test. For analyses with more than 2 comparisons, p-values were adjusted for multiple hypothesis testing using Holm’s method.

Differentially expressed genes (Tables S2 and S3) were determined by bootstrapping, as described in the section “Differential gene expression”. A total of 1,728,000 bootstrap replicates were performed and the resulting p-values were adjusted for multiple hypothesis testing using the Benjamini-Hoschberg procedure (False Discovery Rate).

The trimmed mean (trim = 0.2) and standard error for the number of transcripts expressed per stage were estimated by bootstrapping, with 2000 bootstrap replicates. The number of samples in each stage can be found in Table S1, which contains full sample metadata for both included and excluded samples (i.e. samples that failed to meet the quality control criteria). All samples were distinct measurements; some samples were collected from the same plant or anther, as described in Table S1. In total, a median of 20 samples passed quality control per stage (range: 7, 88), collected from 67 anthers and 25 plants.

### Data availability

Sequencing data are deposited to the Gene Expression Omnibus (accession no. GSE175916) and will be made public after peer review.

## Acknowledgements

We thank J. Ross-Ibarra and E. Josephs for helpful discussions; J. Dinneny for use of the SP8 confocal microscope; and S. Liu for providing the A188 genome sequence ahead of publication. This work was supported by National Science Foundation award 17540974.

## Author Contributions

B.N. and V.W. designed experiments, discussed results, and wrote the manuscript. B.N. conceived the project, performed experiments, and conducted data analysis.

## Competing interests

The authors declare no competing interests.

---

**Figure S1.**
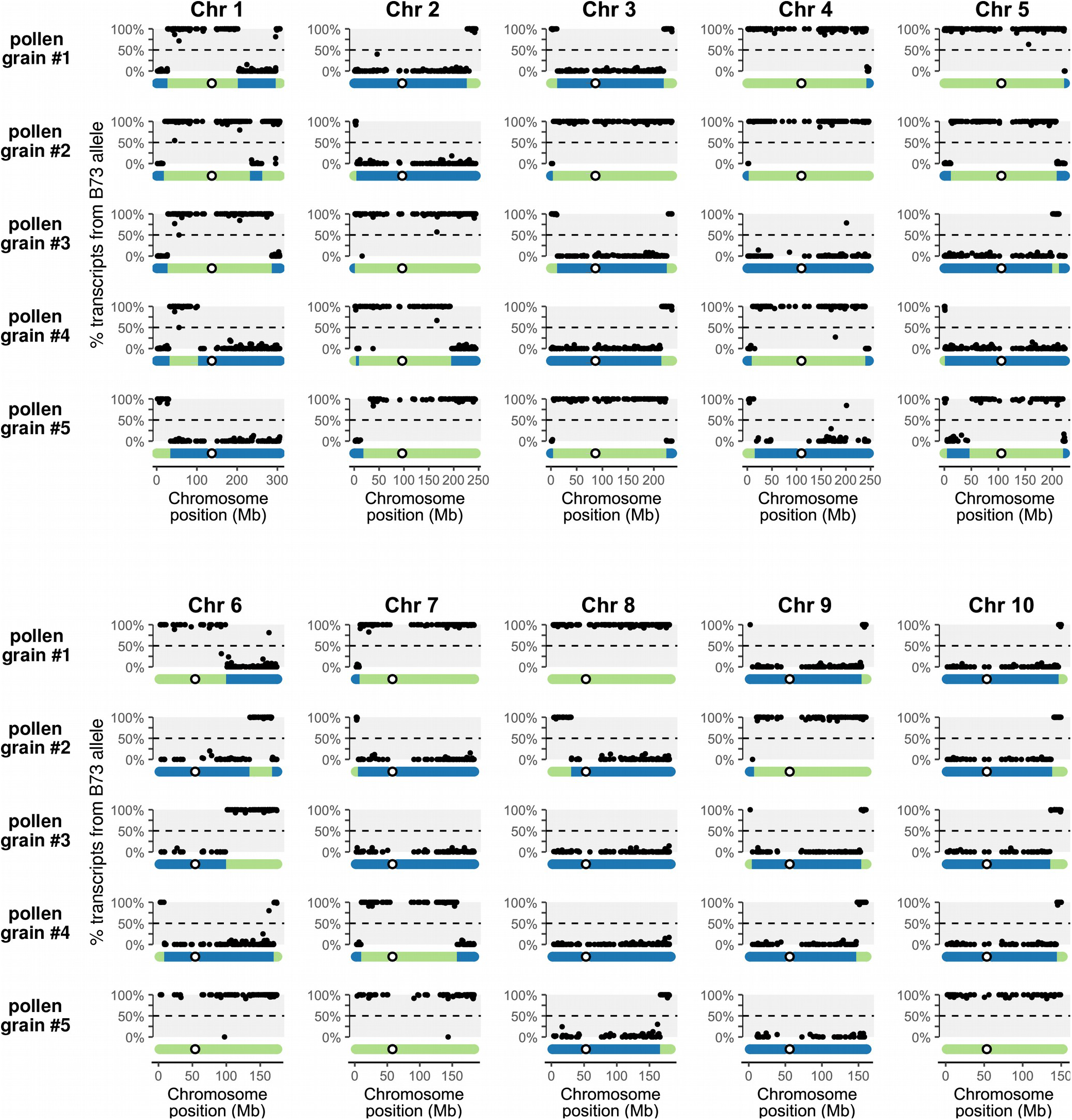
Allelic bias vs chromosomal location for 5 representative pollen grains. Genes with ≥10 genoinformative transcripts are shown. Pollen grain #1 is the same pollen grain analyzed in Fig. 1c,e, here showing the allelic bias across all 10 chromosomes. Inferred haplotypes are depicted below.

**Figure S2.**
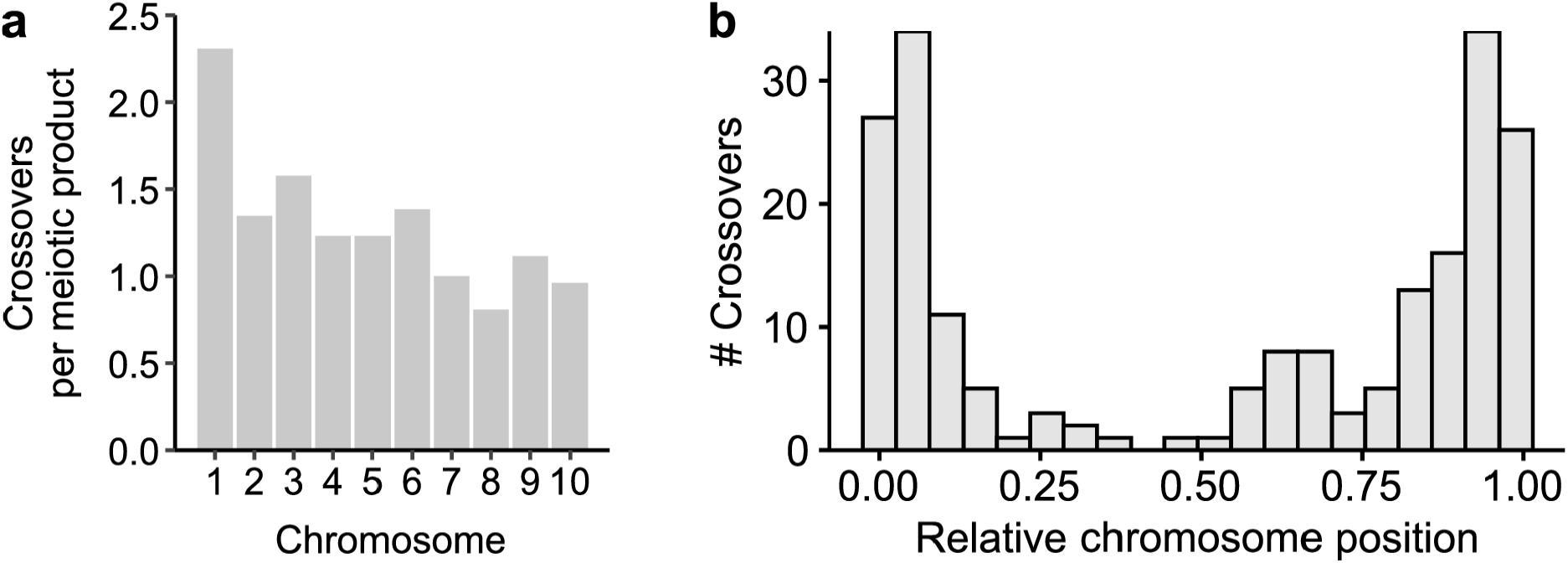
Crossovers inferred from allele-specific RNA-seq data reproduce known characteristics of maize recombination. (**a**) Boxplots showing the number of crossovers inferred for each chromosome per meiotic product (e.g. single pollen grain or microspore). Larger chromosomes had more frequent crossovers, with an overall average of 1.36 crossovers per chromosome. (**b**) Inferred crossovers were more frequent towards chromosome ends, consistent with the known crossover distribution in maize^18^.

**Figure S3.**
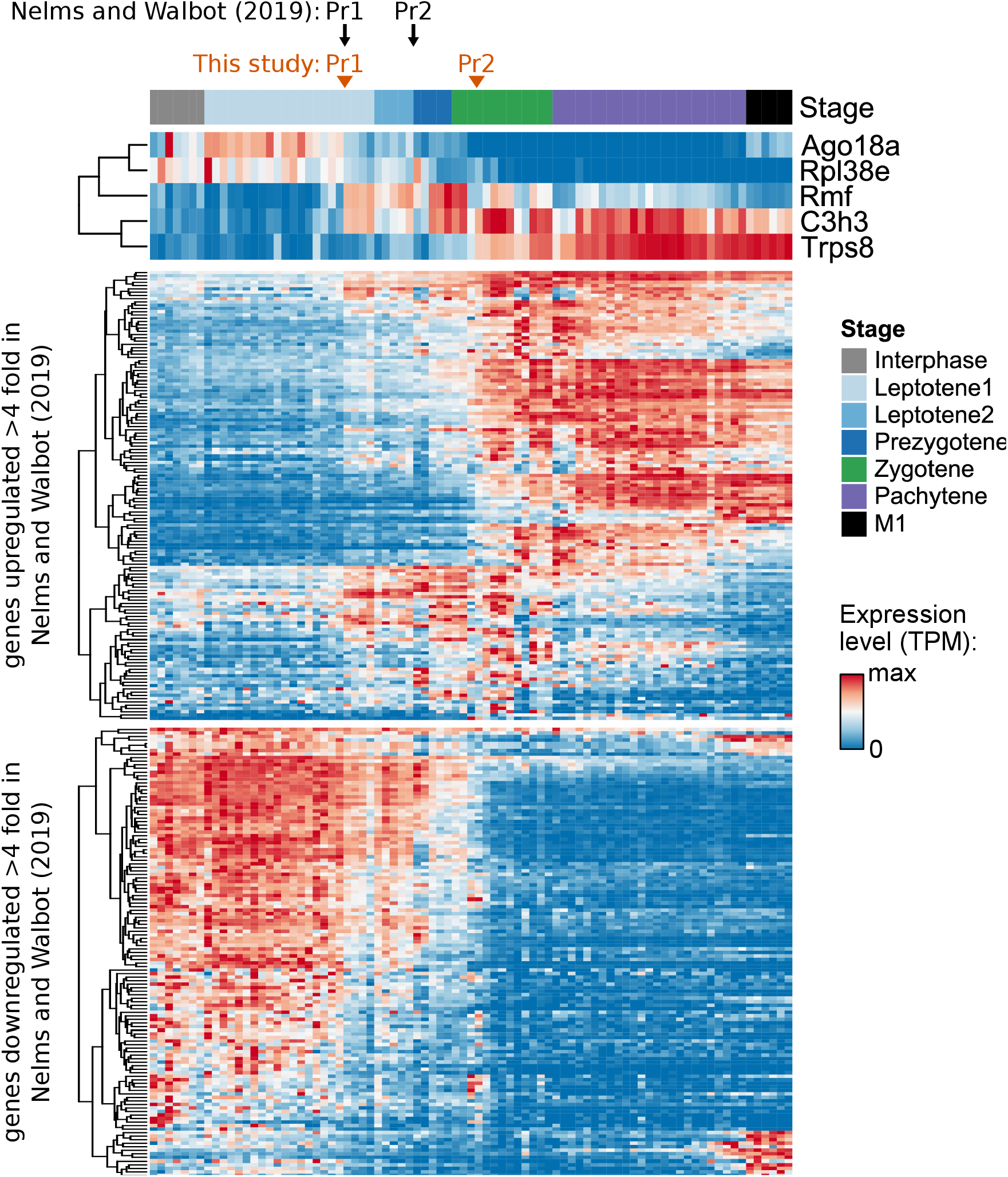
Comparison with Nelms and Walbot (2019). We previously observed a large change in gene expression during meiotic prophase I that took place in two steps^19^, labeled “prophase transition 1” and 2 (Pr1 and Pr2). The heatmaps above show the expression of Pr1/Pr2 genes (identified in our previous report) in the newly obtained data. (Top) Expression of Pr1/Pr2 marker genes. (Middle) Expression of genes up-regulated by 4-fold or more during Pr1/Pr2 in ref 19. (Bottom) Expression of genes down-regulated by 4-fold or more during Pr1/Pr2 in ref 19. Nearly all genes were regulated in a similar direction during meiotic prophase I. However, there were important differences in the timing of the Pr1/Pr2 expression changes compared to ref. 19: (i) a larger number of genes changed in expression during Pr2 here, while more genes changed during Pr1 in our previous report; (ii) Pr2 was slightly later in this dataset, occurring within zygotene. These differences may be the result of variation between maize inbred lines, as our original data were obtained from the W23 inbred line while these data were from an A188 x B73 F1 hybrid.

**Figure S4.**
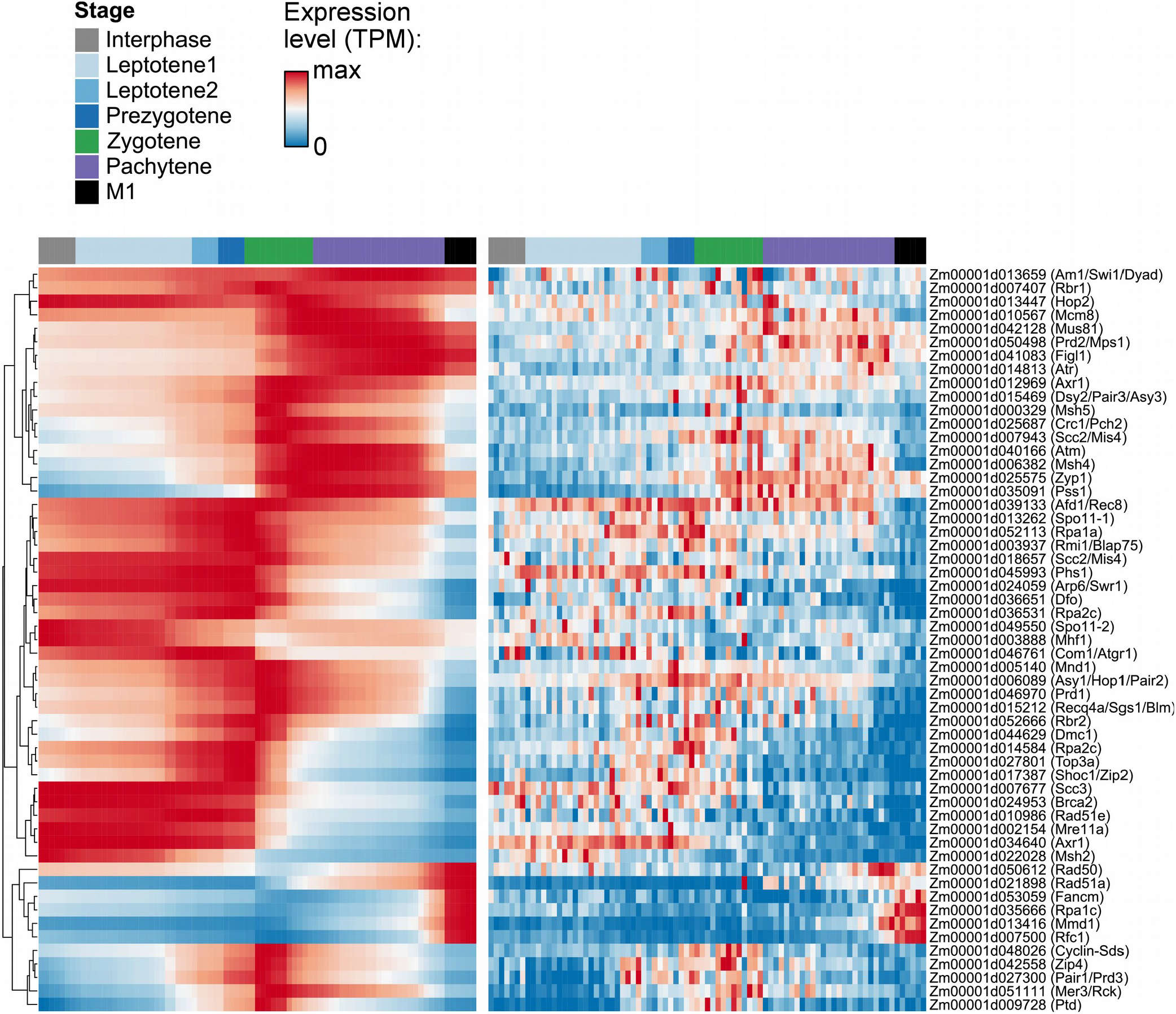
Expression of meiotic genes. Heatmap showing the expression of genes with a putative function in meiosis. The heatmap on the right shows the raw transcript counts while the heatmap on the left was smoothed using a 4^th^-degree polynomial to make it easier to visualize trends. Genes were taken from Table S4 of Nelms and Walbot (2019); genes with established functions in maize meiosis as well as potential orthologs of meiotic genes in other species were evaluated.

**Figure S5.**
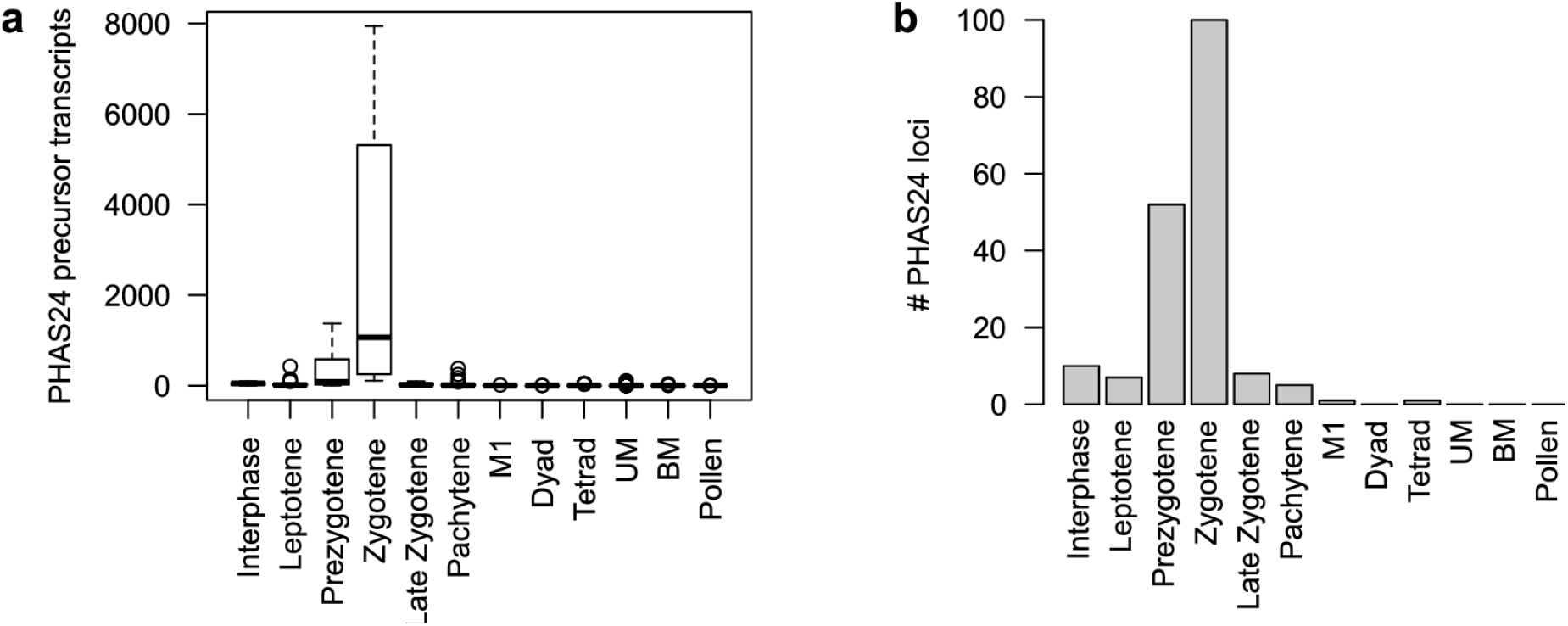
Expression of 24-nucleotide phased small RNA (phasiRNA) precursors. (**a**) Total 24-nt phasiRNA precursor transcripts detected by stage. Boxplots show the median (horizontal line), interquartile range (shaded area; IQR), and whiskers extending up to 1.5 x IQR. (**b**) Number of 24-nt phasiRNA loci expressed at a level of 1 or more transcripts per pollen precursor. PhasiRNA primary transcripts show a burst of expression in early meiotic prophase. A total of 119 phasiRNA loci were considered in this analysis, obtained by remapping previously reported loci^54^ from the v3 and v4 maize genome and excluding any loci that overlap protein-coding genes.

**Figure S6.**
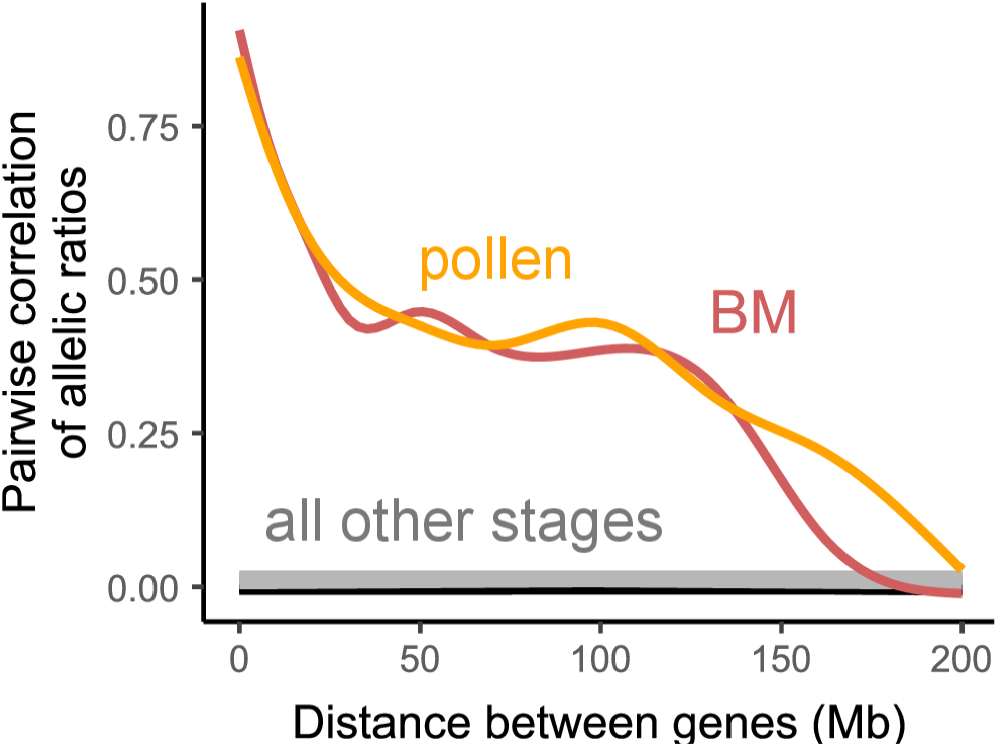
Correlation between allelic ratio and chromosomal distance, by stage. Allelic ratios were calculated as the number of transcripts with the B73 allele divided by the total number of genoinformative transcripts for each gene within each single pollen precursor. Then the Pearson’s correlation was calculated between the allelic ratios for all gene pairs at a given chromosomal distance, and the results were average for precursors at a developmental stage. Genes located near each other on the chromosome were more likely to come from the same parental allele in pollen and BM but not earlier stages. Figure inspired by Bhutani *et al.* (2021).

**Figure S7.**
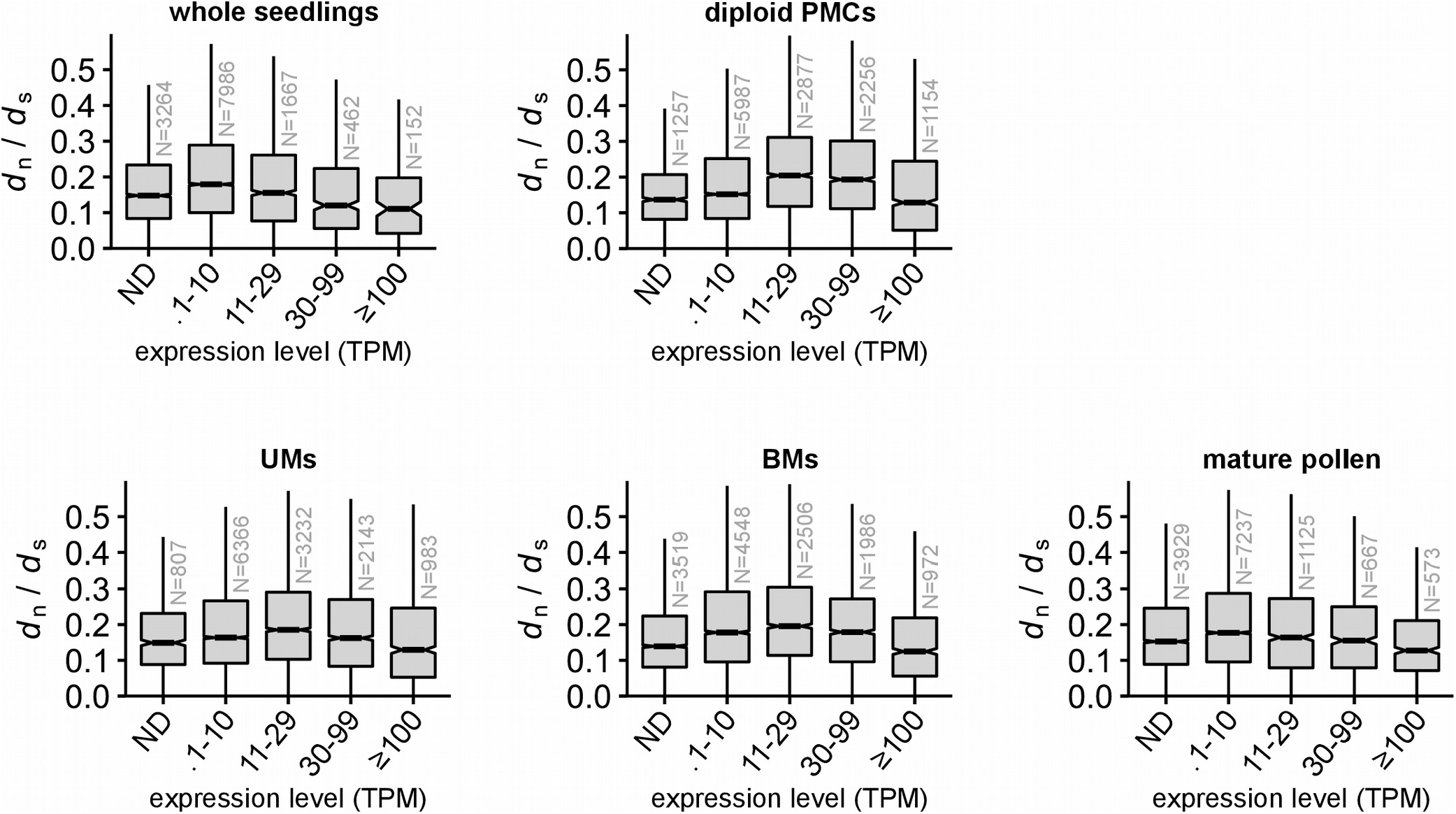
Substitution rate vs expression level in different tissues. The ratio of the number of nonsynonymous substitutions per nonsynonymous site (*d*_n_) to the number of synonymous substitutions per synonymous site (*d*_s_) for genes expressed at different levels in the listed tissues. Diploid tissues are on top and haploid tissues on the bottom. All tissues show a non-monotonic relationship with expression level vs *d*_n_/*d*_s_. Boxplots show the median (horizontal line), interquartile range (shaded area; IQR), and whiskers extending up to 1.5 x IQR. TPM, transcripts per million; ND, not detected; PMCs, pollen mother cells; UMs, unicellular microspores; BMs, bicellular microspores. The reasons *d*_n_/*d*_s_ increases with expression level for low-expressed genes are not clear. One hypothesis is that (i) while purifying selection more often reduces *d*_n_/*d*_s_ for genes with higher expression levels, (ii) transcription-coupled DNA damage leads to a higher mutation rate with increasing transcription; these two mechanisms have opposing effects, and for low-expressed genes the effect of DNA damage outweighs purifying selection while for moderate-expressed genes the balance favors purifying selection. An alternative hypothesis is that, because maize went through a recent whole-genome duplication, the low-expressed genes are more often duplicated genes that are experiencing weakened selection during the process of pseudogenization; genes were excluded from the *d*_n_/*d*_s_ calculations if there was not a clear best match between maize and sorghum (the species used to calculate *d*_n_/*d*_s_), but perhaps the recent gene duplication still has an effect here. While the reasons for the non-monotonic relationship between *d*_n_/*d*_s_ and expression level were not clear, this effect was seen in all tissues and complicates the analysis for low-expressed genes. We focused our main-text analysis on moderately-expressed genes (>100 TPM), a threshold where all tissues show decreasing *d*_n_/*d*_s_ with expression level.

**Figure S8.**
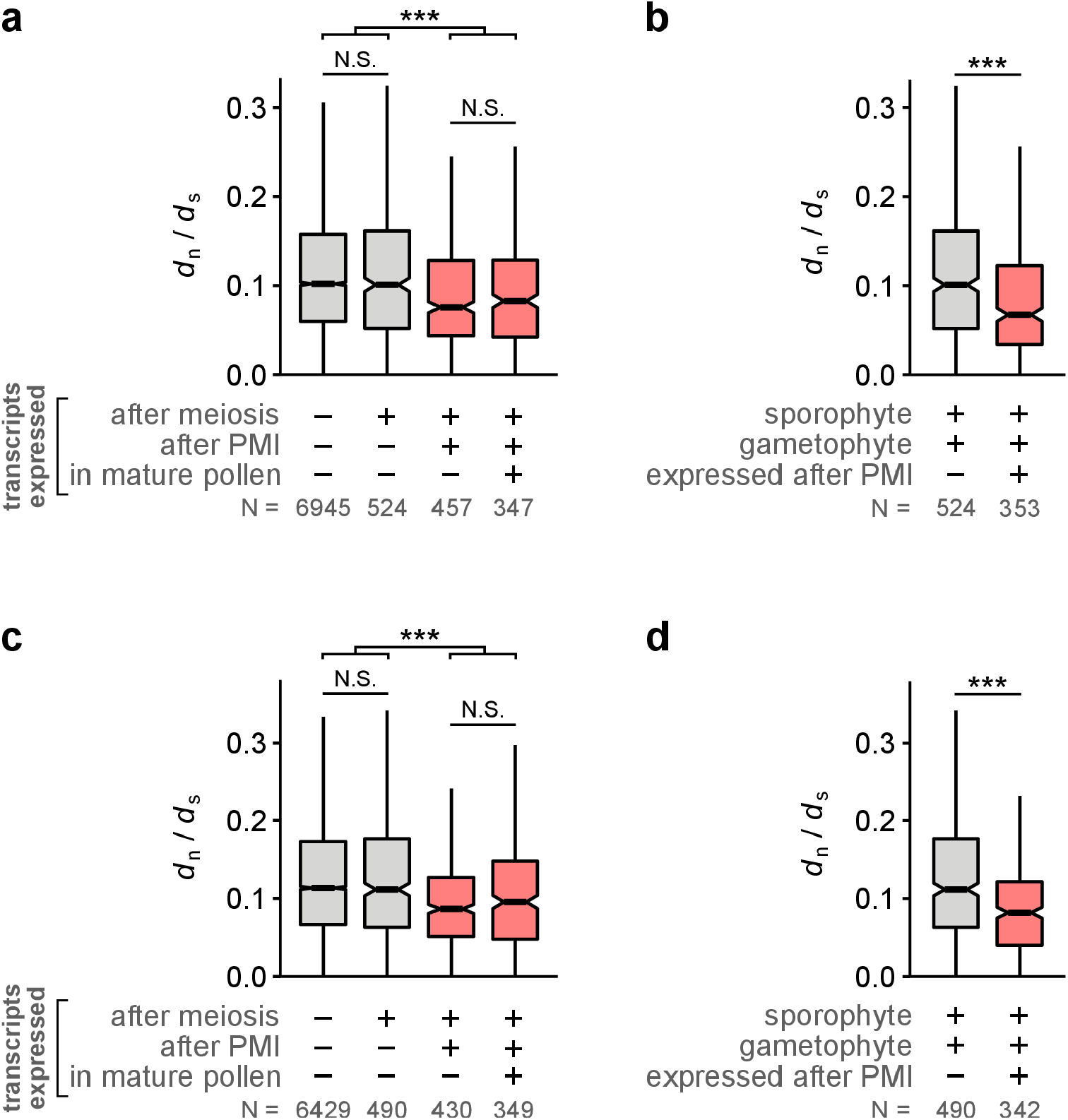
Conservation of gametophyte-expressed genes, in comparison to rice and *Brachypodium*. The ratio of the number of nonsynonymous substitutions per nonsynonymous site (*d*_n_) to the number of synonymous substitutions per synonymous site (*d*_s_) for different gene categories. This figure reproduces the results of Fig. 3a and 3d using different species to calculate *d*_n_/*d*_s_. In Fig. 3, *d*_n_/*d*_s_ was calculated by comparing maize to sorghum, while in this figure it was calculated by comparing maize to rice (panels **a**, **b**) and maize to *Brachypodium distachyon* (panels **c**, **d**). Boxplots show the median (horizontal line), interquartile range (shaded area; IQR), and whiskers extending up to 1.5 x IQR. Gene categories expressed after PMI are shaded red. ***, p < 0.001, Wilcoxon test adjusted for multiple hypothesis testing with Holm’s method.

**Figure S9.**
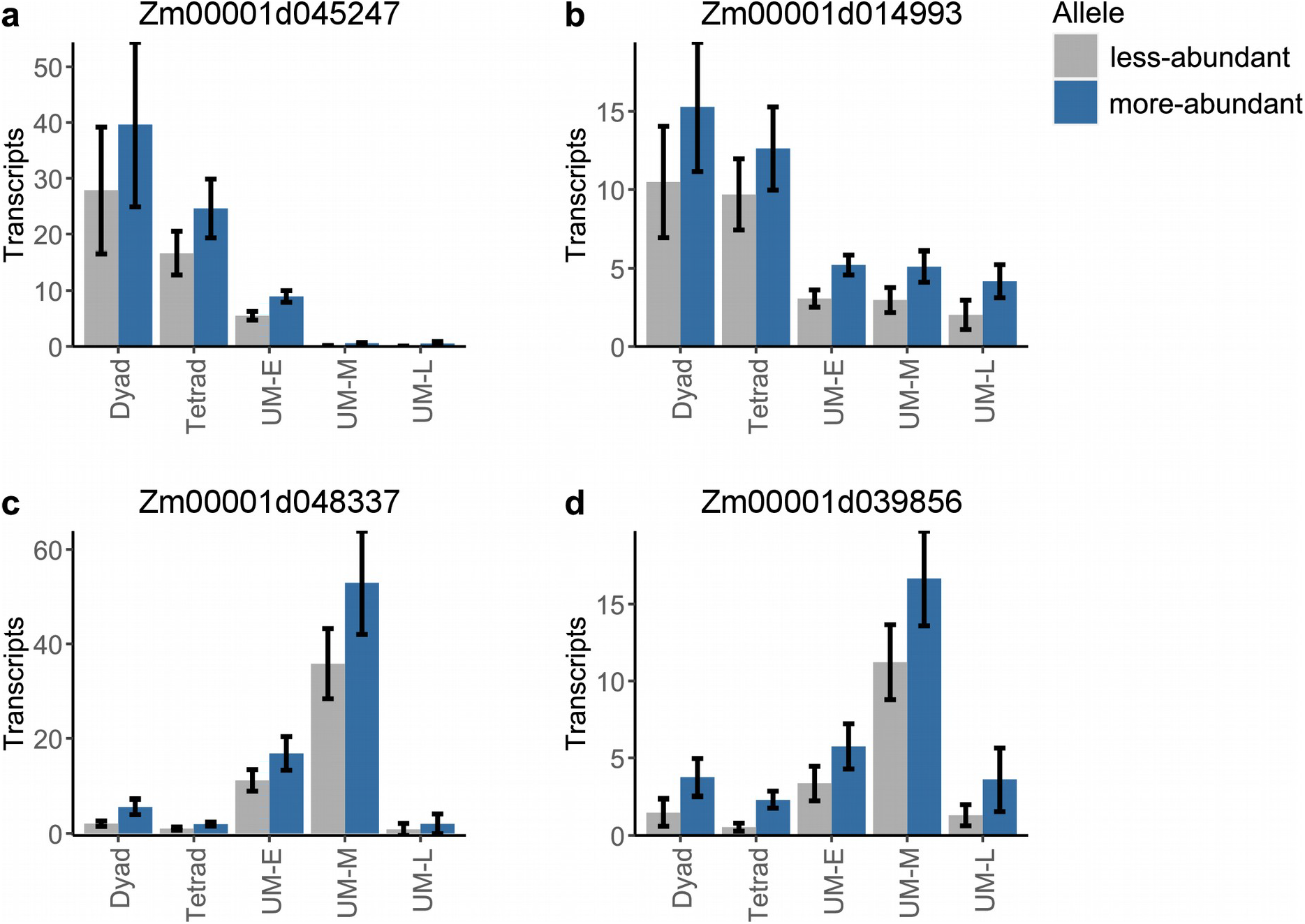
Expression of selected genes from the dyad through UM stages. Gray bars show the average number of transcripts detected for the less-abundant allele while blue bars show the more-abundant allele. Shown are example genes that (**a**) decay in abundance quickly during the UM stage, (**b**) persist during the UM stage, (**c**,**d**) increase in transcript abundance from both alleles during the UM stage. All values are trimmed mean (trim = 0.2) ± standard error, calculated by bootstrapping.

**Table S1.** Metadata for single pollen precursor samples collected for this study.

**Table S2.** Differentially expressed genes during pollen development, focusing on changes in absolute transcript abundance (transcripts per precursor).

**Table S3.** Differentially expressed genes during pollen development, focusing on changes in relative transcript abundance (transcripts per million).

**Table S4.** Sequence of CEL-Seq primers used in this study. The CEL-Seq barcode sequence is underlined.

## Supplementary Note 1

On Illumina patterned flow cells (e.g. Illumina HiSeq 4000, Novaseq) a fraction of multiplexed sample indexes can be exchanged during sequencing, producing incorrect sample assignments^47, 48^. Our libraries were prepared with two sample barcodes: an Illumina index (contained on the i7 Illumina adapter) and an internal CEL-seq barcode contained within the first read of paired-end sequencing:

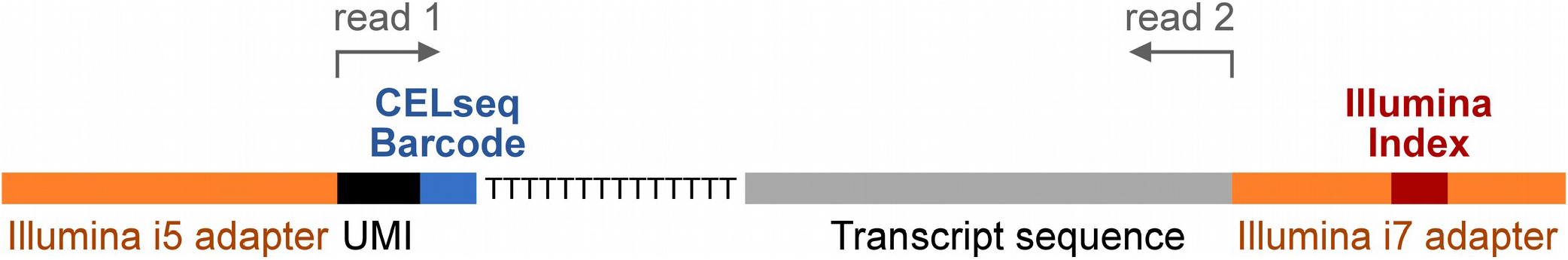

All samples had a unique combination of CELseq barcodes and Illumina indexes, but some samples shared one or the other barcode (for instance, two samples might have the same Illumina index but different CELseq barcodes). To evaluate the index hopping artifact, we compared samples from different stages that either shared or did not share one of the two barcodes. For example, below is a heatmap showing the pairwise Pearson’s correlation between a selection of pollen and pachytene (meiosis) samples. In this heatmap, pollen samples were chosen that share either a CELseq barcode or an Illumina index with at least one pachytene sample. Pachytene samples were divided into 3 groups: those that have no indexes in common with any pollen sample, those that share a CELseq barcode with at least one pollen sample, and those that share an Illumina index with at least one pollen sample:

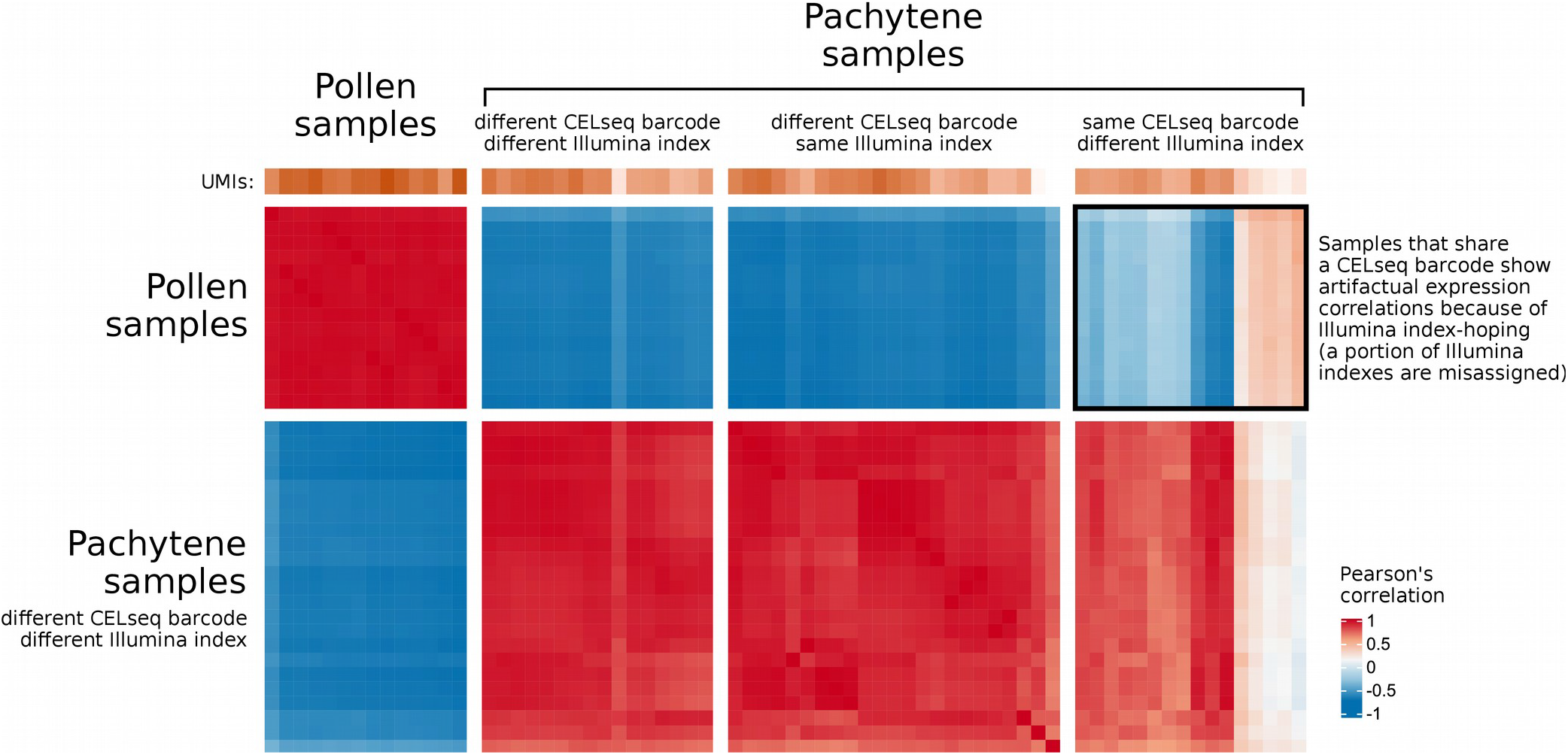

As can be seen above, pachytene and pollen samples that do not share any barcoding information have distinctive transcription profiles and do not correlate with each other (1^st^ and 2^nd^ groups of samples). Samples that share an Illumina index but have different CELseq barcodes also do not show correlation between samples of different types (3^rd^ group of samples); thus, the CELseq barcodes remain reliably associated with the correct transcript. Pachytene and pollen samples that share a CELseq barcode but have different Illumina indexes, however, show correlated expression that is indicative of Illumina index hopping (4^th^ group of samples; see boxed region above).

We also observed sequencing reads mapping to impossible barcode combinations (combinations of CEL-seq and Illumina barcodes that were not used in the experiment); on average, 2.9% of reads for a given Illumina index mapped to impossible barcode combinations, consistent with reported index-hopping levels of 0.5%-10%^29, 30^. Although this amount of index-hopping has been tolerated in prior single-cell RNA-sequencing experiments^48^, we were concerned that this sequencing artifact could affect our results. We took several steps to mitigate the impact of index-hopping on our data.

### Data correction and filtering

First, the data were corrected using a published method^48^ to discard reads that likely arise from index-hopping. Briefly, reads mapping to the same gene with the same CEL-seq barcode and UMI were unlikely to appear in multiple samples by chance, and instead were probably the result of index-hopping; such reads were discarded unless one sample contributed >80% of all instances with a particular CEL-seq barcode, UMI, and gene combination (which would suggest a sample is the sample-of-origin for the transcript). This step reduced the percentage of reads mapping to impossible barcode combinations from 2.9% down to 0.98%.

Second, samples were excluded if there was evidence of contamination by index-hoping based on either of two criteria: 91 sample were excluded because they contributed fewer than 2.4% of all reads matching a particular CEL-seq barcode. Such samples were at greater risk of index-hopping artifacts; a threshold of 2.4% was chosen because it was sufficient to eliminate all samples with impossible barcode combinations. Next, the data were manually examined for evidence of excess correlations that could arise from index-hoping. Heatmaps were constructed showing the pairwise Pearson’s correlation between samples at each pair of developmental stages (similar to the heatmap on the previous page), and samples were removed if they both (i) shared a CEL-seq barcode with samples in the other stage and (ii) showed a greater correlation with the other stage compared to samples that did not have a CEL-seq barcode in common. Neither of these exclusion criteria incorporated information about allele calls, only the total transcript abundance; this was an intentional choice to make sample selection blind to information about biallelic vs monoallelic expression. See Table S1 for a complete list of the excluded samples.

### Validating conclusions with independent RNA-seq data

To confirm our results, we obtained new data under conditions where index-hopping was eliminated. An additional 192 pollen precursors were collected from the UM through BM stages. RNA-sequencing libraries were then prepared with the 96 CELseq barcodes (Table S4) and only one Illumina index. These libraries were sequenced on two separate HiSeq 4000 lanes (2 lanes x 96 indexes = 192 samples). This strategy avoided using the Illumina indexes for sample identification, thus bypassing the index-hopping artifact (the artifact affects the Illumina indexes but not the internal CELseq barcodes, e.g. see heatmap above). These additional data reproduced all findings from our initial lane of sequencing, and all main text figures include data from both the original and additional sequencing lanes.

### Conclusions and recommendations on index-hopping

For the current generation of Illumina sequencers that use patterned flow cells, it is important to be aware of index-hopping to avoid artifactual conclusions. Illumina’s recommendations are to use Unique Dual Indexes, an approach we are migrating towards.

Single-cell data with extensive multiplexing have the potential to be particularly sensitive to index-hopping artifacts; however, it is reassuring that most single-cell data currently available from plants are likely unaffected: our prior single-cell data^19^ was collected using an older HiSeq 2500 instrument, which does not suffer from the index-hopping artifact. The majority of other plant single-cell data have been obtained using the 10X genomics platform. Libraries constructed with 10X are likely free of index-hopping because 10X uses barcodes that are internal to the Illumina adapters (in the same position as the CELseq barcodes used here) and index-hopping specifically affects indexes in the Illumina adapter regions.

## Notes

### Competing Interest Statement

The authors have declared no competing interest.

## References

1. Immler, S. Haploid selection in “diploid” organisms. Annu. Rev. Ecol. Evol. Syst. 50, 1–18 (2019).

2. Mulcahy, D. L., Sari-Gorla, M. & Mulcahy, G. Pollen selection - Past, present and future. Sex. Plant Reprod. 9, 353–356 (1996).

3. Beaudry, F. E. G., Rifkin, J. L., Barrett, S. C. H. & Wright, S. I. Evolutionary genomics of plant gametophytic selection. Plant Commun. 1, 100115 (2020).

4. Tanksley, S., Zamir, D. & Rick, C. Evidence for extensive overlap of sporophytic and gametophytic gene expression in *Lycopersicon esculentum*. Science 213, 453–455 (1981).

5. Khush, G. S. Studies on the linkage map of chromosome 4 of the tomato and on the transmission of induced deficiencies. Genetica 38, 74–94 (1967).

6. Kindiger, B., Beckett, J. B. & Coe, E. H. Differential effects of specific chromosomal deficiencies on the development of the maize pollen grain. Genome 34, 579–594 (1991).

7. Boavida, L. C. et al. A collection of Ds insertional mutants associated with defects in male gametophyte development and function in *Arabidopsis thaliana*. Genetics 181, 1369–1385 (2009).

8. Armbruster, W. S. & Rogers, D. G. Does pollen competition reduce the cost of inbreeding? Am. J. Bot. 91, 1939–1943 (2004).

9. Mulcahy, D. L. Correlation between speed of pollen tube growth and seedling height in *Zea mays L*. Nature 249, 491–493 (1974).

10. Sandler, G., Beaudry, F. E. G., Barrett, S. C. H. & Wright, S. I. The effects of haploid selection on Y chromosome evolution in two closely related dioecious plants. Evol. Lett. 2, 368–377 (2018).

11. Lenormand, T. & Dutheil, J. Recombination difference between sexes: A role for haploid selection. PLoS Biol. 3, e63 (2005).

12. Domínguez, E., Cuartero, J. & Fernández-Muñoz, R. Breeding tomato for pollen tolerance to low temperatures by gametophytic selection. Euphytica 142, 253–263 (2005).

13. Clarke, H. J., Khan, T. N. & Siddique, K. H. M. Pollen selection for chilling tolerance at hybridisation leads to improved chickpea cultivars. Euphytica 139, 65–74 (2004).

14. Warman, C. et al. High expression in maize pollen correlates with genetic contributions to pollen fitness as well as with coordinated transcription from neighboring transposable elements. PLoS Genet. 16, e1008462 (2020).

15. Hsu, S., Huang, Y. & Peterson, P. Development pattern of microspores in *Zea mays L*. - the maturation of upper and lower florets of spikelets among an assortment of genotypes. Maydica 33, 77–98 (1988).

16. Schulz, K. N. & Harrison, M. M. Mechanisms regulating zygotic genome activation. Nat. Rev. Genet. 20, 221–234 (2019).

17. Sidhu, G. K., Fang, C., Olson, M. A., Falque, M. & Martin, O. C. Recombination patterns in maize reveal limits to crossover homeostasis. Proc. Natl. Acad. Sci. 112, (2015).

18. Kianian, P. M. A. et al. High-resolution crossover mapping reveals similarities and differences of male and female recombination in maize. Nat. Commun. (2018). doi:10.1038/s41467-018-04562-5

19. Nelms, B. & Walbot, V. Defining the developmental program leading to meiosis in maize. Science 364, 52–56 (2019).

20. Gossmann, T. I., Schmid, M. W., Grossniklaus, U. & Schmid, K. J. Selection-driven evolution of sex-biased genes is consistent with sexual selection in *Arabidopsis thaliana*. Mol. Biol. Evol. 31, 574–583 (2014).

21. Arunkumar, R., Josephs, E. B., Williamson, R. J. & Wright, S. I. Pollen-specific, but not sperm-specific, genes show stronger purifying selection and higher rates of positive selection than sporophytic genes in *Capsella grandiflora*. Mol. Biol. Evol. 30, 2475–2486 (2013).

22. Chettoor, A. M. et al. Discovery of novel transcripts and gametophytic functions via RNA-seq analysis of maize gametophytic transcriptomes. Genome Biol. 15, 1–23 (2014).

23. Twell, D., Oh, S. A. & Honys, D. Pollen development, a genetic and transcriptomic view. in The Pollen Tube. Plant Cell Monographs (ed. Malhó, R.) 3, 15–45 (2006).

24. Honkela, A. et al. Genome-wide modeling of transcription kinetics reveals patterns of RNA production delays. Proc. Natl. Acad. Sci. U. S. A. 112, 13115–13120 (2015).

25. Jia, H., Suzuki, M. & Mccarty, D. R. Regulation of the seed to seedling developmental phase transition by the LAFL and VAL transcription factor networks. Wiley Interdiscip. Rev. Dev. Biol. 3, 135–145 (2014).

26. Valdivia, E. R., Sampedro, J., Lamb, J. C., Chopra, S. & Cosgrove, D. J. Recent proliferation and translocation of pollen group 1 allergen genes in the maize genome. Plant Physiol. 143, 1269–1281 (2007).

27. Stelpflug, S. C. et al. An expanded maize gene expression atlas based on RNA-sequencing and its use to explore root development. Plant Genome 9, (2016).

28. Harris, B. & Dure, L. I. Developmental regulation in cotton seed germination: Polyadenylation of stored messenger RNA. Biochemistry 17, 3250–3256 (1978).

29. Kim, N. & Jinks-Robertson, S. Transcription as a source of genome instability. Nat. Rev. Genet. 13, 204– 214 (2012).

30. McCormick, S. Male gametophyte development. Plant Cell 5, (1993).

31. Eady, C., Lindsey, K. & Twell, D. The significance of microspore division and division symmetry for vegetative cell-specific transcription and generative cell differentiation. Plant Cell 7, 65–74 (1995).

32. Zhang, J. et al. Sperm cells are passive cargo of the pollen tube in plant fertilization. Nat. Plants 3, 1–5 (2017).

33. Glöckle, B. et al. Pollen differentiation as well as pollen tube guidance and discharge are independent of the presence of gametes. Dev. 145, (2018).

34. Bedinger, P. A. & Edgerton, M. D. Developmental staging of maize microspores reveals a transition in developing microspore proteins. Plant Physiol. 92, 474–479 (1990).

35. Slotkin, R. K. et al. Epigenetic reprogramming and small RNA silencing of transposable elements in pollen. Cell 136, 461–472 (2009).

36. Borg, M. & Berger, F. Chromatin remodelling during male gametophyte development. Plant J. 83, 177– 188 (2015).

37. Bhutani, K. et al. Widespread haploid-biased gene expression enables sperm-level natural selection. Science 371, eabb1723 (2021).

38. Dawe, R. K., Sedat, J. W., Agard, D. A. & Cande, W. Z. Meiotic chromosome pairing in maize is associated with a novel chromatin organization. Cell 76, 901–912 (1994).

39. Hashimshony, T. et al. CEL-Seq2: sensitive highly-multiplexed single-cell RNA-Seq. Genome Biol. 17, 77 (2016).

40. Kivioja, T. et al. Counting absolute numbers of molecules using unique molecular identifiers. Nat. Methods 9, 72–74 (2011).

41. Chen, S., Zhou, Y., Chen, Y. & Gu, J. Fastp: An ultra-fast all-in-one FASTQ preprocessor. Bioinformatics 34, i884–i890 (2018).

42. Jiao, Y. et al. Improved maize reference genome with single-molecule technologies. Nature 546, 524–527 (2017).

43. Lin, G. et al. Chromosome-level genome assembly of a regenerable maize inbred line A188. Genome Biol. 22, 1–30 (2021).

44. Pertea, M., Kim, D., Pertea, G. M., Leek, J. T. & Salzberg, S. L. Transcript-level expression analysis of RNA-seq experiments with HISAT, StringTie and Ballgown. Nat Protoc. 11, 1650–1667 (2016).

45. Krueger, F. & Andrews, S. R. SNPsplit: Allele-specific splitting of alignments between genomes with known SNP genotypes. F1000Research 5, 1–16 (2016).

46. Smith, T., Heger, A. & Sudbery, I. UMI-tools: Modelling sequencing errors in Unique Molecular Identifiers to improve quantification accuracy. Genome Res. (2017).

47. Sinha, R. et al. Index switching causes “spreading-of-signal” among multiplexed samples in Illumina HiSeq 4000 DNA sequencing. bioRxiv 125724 (2017). doi:10.1101/125724

48. Griffiths, J. A., Richard, A. C., Bach, K., Lun, A. T. L. & Marioni, J. C. Detection and removal of barcode swapping in single-cell RNA-seq data. Nat. Commun. 9, 1–6 (2018).

49. Trapnell, C. et al. The dynamics and regulators of cell fate decisions are revealed by pseudotemporal ordering of single cells. Nat. Biotechnol. 32, 381–386 (2014).

50. Hastie, T. & Stuetzle, W. Principal curves. J. Am. Stat. Assoc. 84, 502–516 (1989).

51. Gu, Z., Eils, R. & Schlesner, M. Complex heatmaps reveal patterns and correlations in multidimensional genomic data. Bioinformatics 32, 2847–2849 (2016).

52. Lyons, E. & Freeling, M. How to usefully compare homologous plant genes and chromosomes as DNA sequences. Plant J. 53, 661–673 (2008).

53. Bailey, T. L. et al. MEME Suite: Tools for motif discovery and searching. Nucleic Acids Res. 37, 202–208 (2009).

54. Zhai, J. et al. Spatiotemporally dynamic, cell-type-dependent premeiotic and meiotic phasiRNAs in maize anthers. Proc. Natl. Acad. Sci. U. S. A. 112, 3146–3151 (2015).

